# Leveraging Scaffold Information to Predict Protein-ligand Binding Affinity with an Empirical Graph Neural Network

**DOI:** 10.1101/2022.08.19.504617

**Authors:** Chunqiu Xia, Shi-Hao Feng, Ying Xia, Xiaoyong Pan, Hong-Bin Shen

## Abstract

**Motivation:** Protein-ligand binding affinity prediction is an important task in structural bioinformatics for drug discovery and design. Although various scoring functions have been proposed, it remains challenging to accurately evaluate the binding affinity of a protein-ligand complex with known bound structure due to the potential preference of scoring system. In recent years, deep learning techniques have been applied to scoring functions without sophisticated feature engineering. Nevertheless, existing methods cannot model the differential contribution of atoms in various regions of proteins, and the relationship between atom properties and intermolecular distance is also not fully explored.

**Results:** We propose a novel empirical graph neural network for accurate protein-ligand binding affinity prediction (EGNA). Graphs of protein, ligand and their interactions are constructed based on different regions of each bound complex. Proteins and ligands are effectively represented by graph convolutional layers, enabling the EGNA to capture interaction patterns precisely by simulating empirical scoring functions. The contributions of different factors on binding affinity can thus be transparently investigated. EGNA is compared with the state-of-the-art machine learning-based scoring functions on two widely used benchmark datasets. The results demonstrate the superiority of EGNA and its good generalization capability.

**Availability and implementation:** The web server and source code of EGNA is available at www.csbio.sjtu.edu.cn/bioinf/EGNA and https://github.com/chunqiux/EGNA.

**Supplementary information:** Supplementary data are available at Bioinformatics online.

## 1. Introduction

Protein-ligand binding affinity measures the strength of the interaction between a protein and a ligand molecule (e.g., protein, nucleotide or drug-like molecules) (Gilson and Zhou, 2007). Accurate binding affinity can improve the understanding of biological processes involving protein-ligand interactions (Furuhashi and Hotamisligil, 2008; Seo, et al., 2014). Furthermore, binding affinity can also be used to rank candidate drugs against a target protein in the drug discovery (Chaires, 2008; McInnes, 2007). However, given a protein structure, it is still time-consuming and labor-intensive to determine its binding affinity to the ligands by wet-lab experiments (e.g., isothermal titration calorimetry, surface plasmon resonance and fluorescence-based methods). Thus, various structure-based computational methods have been proposed for protein-ligand binding affinity prediction.

One typical protein-ligand binding affinity prediction method is based on free energy simulations, such as free energy perturbation and thermodynamic integration. It integrates molecular dynamics simulation with free energy sampling algorithms and is considered as one of the most rigorous computational methods (Cournia, et al., 2020). In general, these methods can provide accurate binding affinity prediction but would suffer from high computational cost (Brandsdal, et al., 2003).

Another popular protein-ligand binding affinity prediction method is based on Scoring Functions (SFs). Compared with free energy-based simulations, scoring functions are used to predict the binding affinity based on the bound structures instead of simulating the entire binding process. Thus, they are much faster and more appropriate in the large-scale binding affinity prediction applications.

Classical scoring functions can be generally divided into three types: physics-based, knowledge-based and empirical scoring functions (Li, et al., 2019). Physics-based scoring functions combine different energy terms derived from force field (Meng, et al., 1992), solvation models (Jorgensen, et al., 1983), quantum mechanics (Raha, et al., 2007), and etc.; Knowledge-based scoring functions are proposed based on the inverse Boltzmann analysis and require the occurrence frequency of any contact atom pair and the same atom pair in a reference state (DeWitte and Shakhnovich, 1996); Empirical scoring functions are generally weighted sum of various energy terms and the weights can be obtained by linear regression (Bohm, 1992) or other algorithms. They are computationally efficient due to their straightforward forms. Nevertheless, their performance still has much space to improve since intermolecular interaction mechanisms are complicated and it remains challenging to approximate the occurrence frequency in a reference state precisely (Liu and Wang, 2015).

Machine learning-based scoring functions, also known as descriptor-based scoring functions, have attracted more attentions in the past decade due to their superior performance (Deng, et al., 2004). Classical SFs use linear combinations of energy terms. Differently, machine learning-based SFs apply nonlinear learning models. For example, RF-Score utilizes a random forest (RF) which is composed of multiple CART trees for regression (Ballester and Mitchell, 2010); AGL-Score integrates algebraic graphs with gradient boosting trees (GBTs) (Nguyen and Wei, 2019); NN-score applies a fully-connected neural network with two hidden layers to predict whether the *K*_d_ value of a protein-ligand complex is less than 25μM (Durrant and McCammon, 2010). Although nonlinear models substantially improve the prediction performance, machine learning-based SFs heavily rely on the feature engineering and expert knowledge.

In recent years, deep learning has achieved remarkable successes in the fields of computer vision (He, et al., 2016), natural language processing (Vaswani, et al., 2017), speech recognition (Deng, et al., 2013), and bioinformatics (Alipanahi, et al., 2015; Xia, et al., 2022) due to its representation capabilities (Jumper, et al., 2021; LeCun, et al., 2015). It has also been applied to protein-ligand binding affinity prediction (Wang, et al., 2021). Unlike conventional machine learning-based methods, deep learning-based scoring functions can directly use raw data, including atom properties, bond information and geometric features, instead of hand-crafted features. Some deep learning-based SFs voxelize complex structures and then apply three-dimensional (3D) convolutional neural networks (CNNs), e.g., K_DEEP_ and Pafnucy (Jimenez, et al., 2018; Stepniewska-Dziubinska, et al., 2018). Other SFs extract occurrence frequencies features of atom pairs like knowledge-based SFs and organize these features as matrices (Seo, et al., 2021; Zheng, et al., 2019). Then, 1D or 2D CNNs are applied to them. Moreover, graph neural networks (GNNs) have also been proved effective for binding affinity prediction by transforming structures to graphs based on bonds or distance between atoms (Li, et al., 2021). Different from 3D CNNs, GNNs are proposed for processing non-Euclidean data and invariant to rotation (Wu, et al., 2021).

In the abovementioned deep learning-based SFs, atoms in the binding pockets or other regions of the protein are treated equally, which may actually have different contribution to the binding affinity. Moreover, the relationship between intermolecular distance and atom properties are not fully explored. These are potential reasons for that the performance of existing deep learning-based SFs does not show much superiority to those conventional machine learning models with sophisticated feature engineering. In order to further improve the deep learning-based SFs, here we propose a novel <u>E</u>mpirical <u>G</u>raph Neural <u>N</u>etwork for accurate protein-ligand binding <u>A</u>ffinity prediction (EGNA). The inputs of our model are bound protein-ligand complexes and different graphs of protein, ligand and their interactions are constructed based on the intermolecular distance. In this model, proteins and ligands are represented separately in EGNA. Inspired by physics-based and empirical SFs, an empirical interaction representation layer is designed to represent the interactions and exchange information between a protein and a ligand, which could further exploit the distance information and raw atom (or amino acid) features. Our method is compared to the state-of-the-art scoring functions on two widely used benchmark datasets (Dunbar, et al., 2011; Su, et al., 2019). The results show that our method EGNA achieves lower errors and higher correlations, demonstrating the efficacy of the new EGNA model.

## 2. Materials and Methods

### 2.1. Benchmark datasets

PDBbind v.2016 is one of the widely used benchmark datasets for evaluating protein-ligand binding affinity prediction (Liu, et al., 2017). It has three subsets: a general set, a refined set and a core set. The general set contains 16,179 protein-small ligand complexes with experimentally determined binding affinity (i.e., *K*_d_, *K*_i_ or *IC*_50_). The refined set is a high-quality subset of the general set and contains 4,057 entries. The core set is a subset of the refined set and contains 285 entries. It is used to evaluate the performance of SFs in the Comparative Assessment of Scoring Functions (CASF) project, denoted as CASF-2016 (Su, et al., 2019). The overlapping entries between the CASF-2016 and the general/refined set are all removed from the latter ones. Then, the general set and refined set is used to train our model while CASF-2016 is used as an independent test set in this study.

The Community Structure-Activity Resource (CSAR) is another dataset constructed for protein-ligand docking and scoring evaluation (Dunbar, et al., 2011). The high-quality version of CSAR (CSAR-HiQ) is composed of two sets, which contain 176 and 167 entries, respectively. We combine the two sets and remove the overlapping entries between the combined set and PDBbind v.2016. The final set contains 49 entries and is used for independent evaluation. The PDB IDs of the 49 protein-ligand complexes can be referred to section A in S1 Supporting Information.

### 2.2. Graph construction and node feature extraction

In this section, we introduce how to prepare the data for our GNN. At first, graphs of proteins, ligands and interactions are constructed from protein-ligand complexes with known bound structures. Then, raw features for each node in protein and ligand graphs are extracted.

#### 2.2.1. Protein, ligand and interaction graph construction

It has been proven feasible to represent the 3D structures of proteins or other molecules as graphs in protein-protein interface identification, protein-ligand binding site prediction and binding affinity prediction (Fout, et al., 2017; Li, et al., 2021; Xia, et al., 2021). As shown Fig 1, we construct a protein graph 𝒢_P_ = (*V*_P_, *E*_P_), a ligand graph 𝒢_L_ = (*V*_L_, *E*_L_), and an interaction graph 𝒢_I_ = (*V*_I_, *E*_I_) for each complex. All the graphs are undirected.

**Fig 1.**
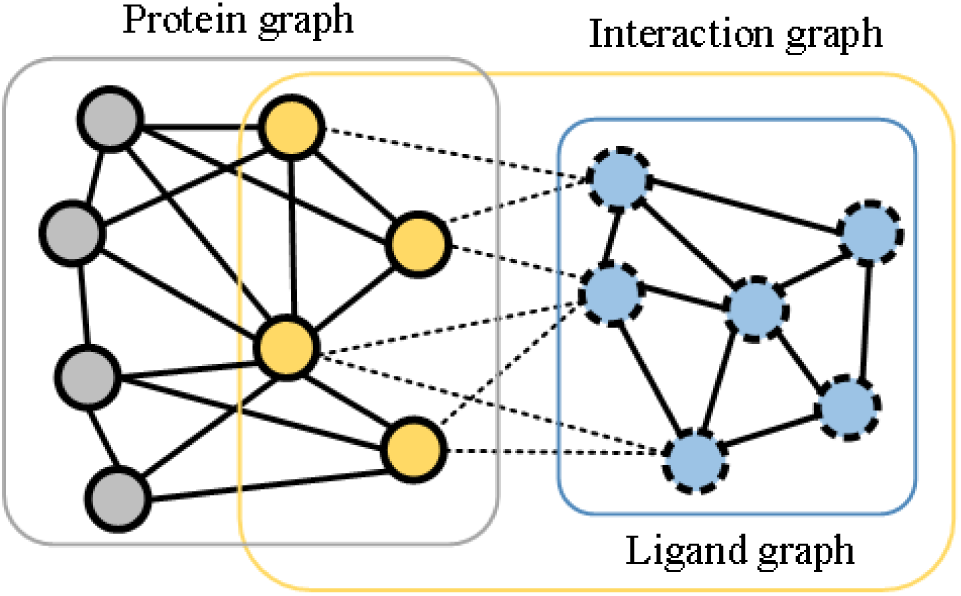
Illustration of a protein graph 𝒢_P_ (grey), a ligand graph 𝒢_L_ (blue) and an interaction graph 𝒢_I_ (yellow) constructed from the of a protein-ligand complex structure. In the interaction graph, only dotted lines represent the edges.

**Fig 2.**
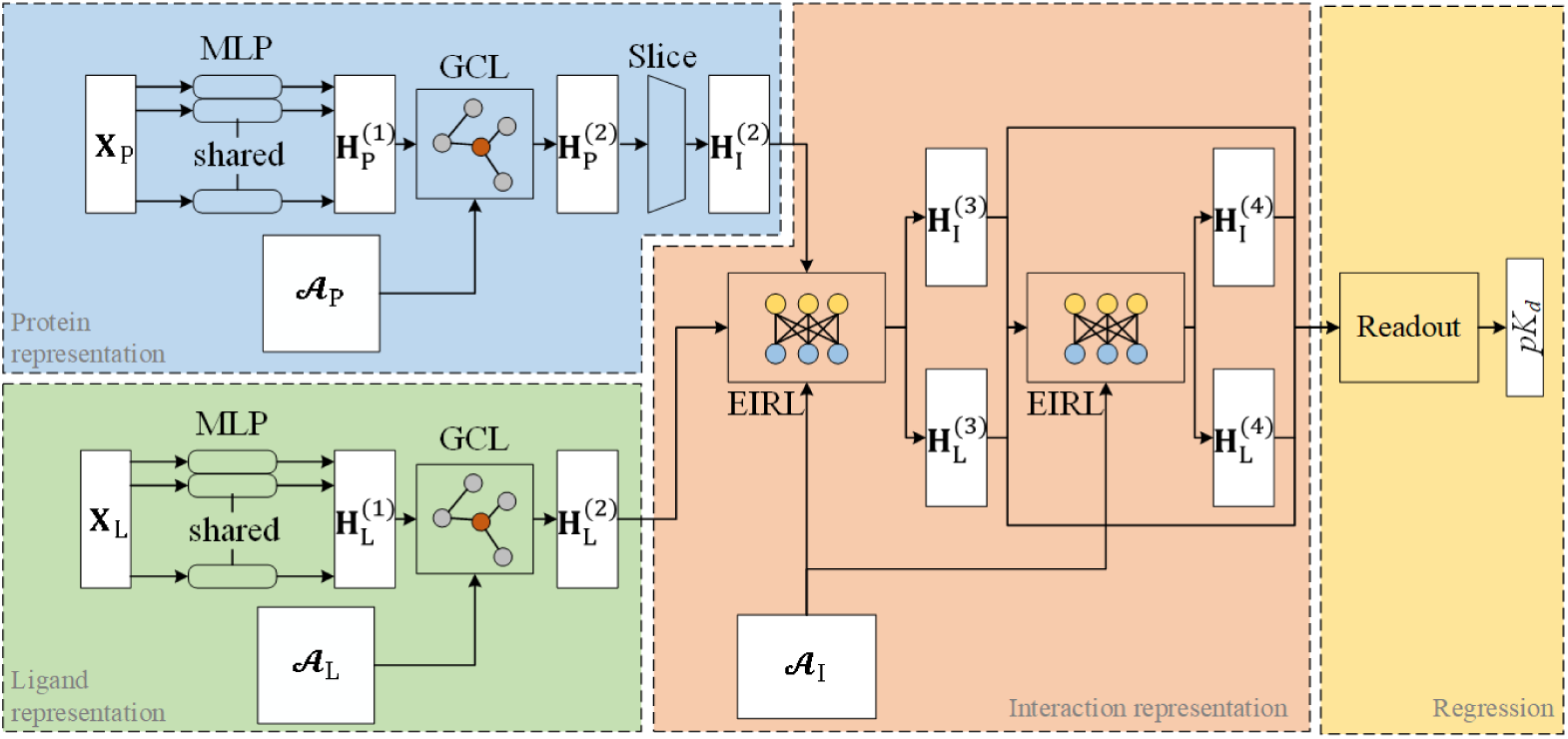
The architecture of the proposed empirical graph neural network. It consists of four modules: protein representation, ligand representation, interaction representation and regression. In the protein/ligand representation module, graph convolutional layers (GCLs) are used to learn node embeddings for molecules. In the interaction representation layer, empirical interaction representation layers (EIRLs) are used to exchange and integrate information from proteins and ligands based on the pairwise distance. In the regression module, the learned node embeddings are aggregated to predict the p*K*_*d*_ value of the protein-ligand complex.

##### Ligand graphs

Each heavy atom in the ligand is considered as a member of the node set *V*_L_. If a bond exists between any two atoms, an edge is defined between the two corresponding nodes. The adjacency matrix of a ligand graph is defined based on *E*_L_ as follows:

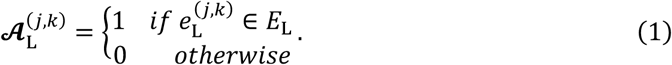

##### Protein graphs

Different from ligand graphs, only a part of the protein structure which is close to the ligand is used to construct the graph. The node set of the protein graph at the residue level is defined as follows:

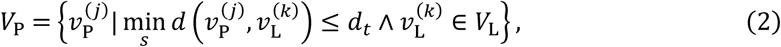

where 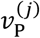 is the *j*^th^ residue of the protein, 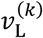 is the *k*^th^ heavy atom of the ligand, and 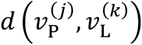 is the Euclidean distance between the *C*_*β*_ atom (*C*_*α*_ for glycine) of 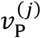 and 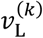.

The distance threshold *d*_*t*_ is set to 20Å in this study according to our experiments.

Protein graphs are complete graphs where any two nodes are connected by an edge. The adjacency matrix of a protein graph is dense and defined based on the distance between two *C*_*β*_ atoms as follows:

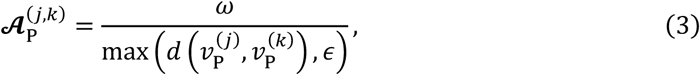

where two hyperparameters *ϵ* > 0 and ω > 0 are used for normalization. Therefore,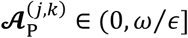..

##### Interaction graphs

𝒢_I_ contains only the residues from the binding pocket of the protein and heavy atoms from the ligand. Its node set is 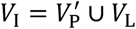. The difference between 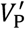 and *V*_P_ is that *d*_*t*_ of the former one is set to 10Å (Liu, et al., 2017), and thus 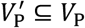. The edge set *E*_I_ is defined based on 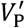 and *V*_L_ as follows:

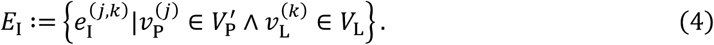

Evidently, 𝒢_I_ is a bipartite graph. Its adjacency matrix 𝒜_I_ is also defined based on the distance between nodes belonging to 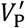 and *V*_L_ similar to Eq.3, respectively.

#### 2.2.2. Node feature extraction

After graph construction, properties related to binding affinity of residues or atoms are extracted. For proteins, we run a hidden Markov model-based multiple sequence alignment (MSA) tool, HHblits, against Uniclust30 to generate sequence profiles for each protein chain (Mirdita, et al., 2017; Remmert, et al., 2011). The sequence profiles contain the emission frequencies, transition frequencies, and MSA diversity of residues in protein sequences. They reflect the tendency of a residue to mutate into other types of residues. Here, features derived from sequence profiles are named as evolutionary features. The procedure of featurization is described in Section B of S1 Supporting Information. It should be noted that only the feature vectors of the residues in *V*_P_ are considered and the length of the feature vector is *m*_P_ = 30.

For ligands, we extract atom-level raw features for each atom in *V*_L_ by OpenBabel (O’Boyle, et al., 2011). Each atom is classified into one of 9 classes according to its atom type (i.e., B, C, N, O, P, S, Se, halogen, and metal) and encoded as a 9-D one-hot vector. We also investigate whether the atom is in ring, aromatic and a hydrogen bond acceptor. These properties are then binarized as a 3-D one-hot vector. Moreover, the partial charge and hybridization of atoms are also encoded as 2-D vector. All these features are concatenated and the length of the feature vectors is *m*_L_ = 14.

### 2.3. Empirical graph neural network

Inspired by physics-based and empirical scoring functions, we propose a novel empirical graph neural network to improve the accuracy of the protein-ligand binding affinity prediction. At first, the architecture of the proposed GNN is overviewed. Then, protein, ligand and interaction representation layers are introduced in detail.

#### 2.3.1. Overall pipeline

Our model EGNA is composed of four modules: a protein representation module, a ligand representation module, an interaction representation module and a regression module. In the first step, a graph convolutional layer (GCL) is used to learn the embeddings of residues/atoms from their neighbors only in the protein/ligand. In the second step, empirical interaction representation layers (EIRLs) are designed to exchange and integrate information from proteins and ligands based on the pairwise distance. At last, the learned node embeddings after the interaction representation layers are aggregated to predict the p*K*_*d*_ of the protein-ligand complex.

In the training stage, we minimize the root mean squared error (RMSE) between the true and predicted p*K*_*d*_ values. Furthermore, a simple bagging strategy is applied for the robustness, where the final output is an average of the predicted p*K*_*d*_ values from multiple independently trained models. The implementation details can refer to the section C in S1 Supporting Information.

#### 2.3.2. Protein and ligand representation

To explore the interactions between proteins and ligands, it is necessary for the network to learn discriminative embeddings of each residue or atom from the raw features. The raw node feature matrices of a protein and a ligand are denoted as 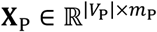 and 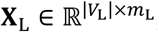, respectively. For a protein, a multi-layer perceptron (MLP) is used to get the initial embeddings 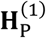 of residues from **X**_P_. Then, a graph convolutional layer is used to aggregate information into the target residue from its spatial neighboring residues as follows:

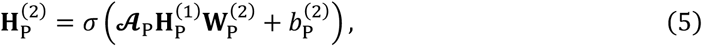

where 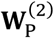 is a learnable weight matrix, 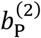 is a learnable bias scalar, and σ(·) is the activation function.

Considering that the binding affinity is more related to the residues in the binding pocket, a slice operation is performed to extract the embeddings of those residues in 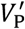, which is denoted as 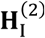. For a ligand, each heavy atom is represented similar to the residues in proteins, where similar MLP and GCL layers are used to learn the embeddings 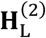 of atoms in the ligand, but with different learnable parameters and their own adjacency matrices 𝒜_L_.

#### 2.3.3. Empirical interaction representation

According to knowledge-based SFs, the binding affinity is highly correlated to the distance patterns of the bimolecular interactions (Schulz-Gasch and Stahl, 2004). Thus, the interaction between proteins and ligands is modeling by GCLs with the dense adjacency matrix 𝒜_I_. If the hyperparameter *ϵ* in Eq.3 is set to a small value, which can guarantee the pairwise distance 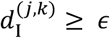 in most cases, each element of 𝒜 will approximately equal to 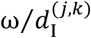. In this study, *ϵ* is set to 2Å and the distance of most residue-atom pairs is larger than it in the training set. The *j*^th^ residue in the protein and the *k*^th^ atom in the ligand can exchange their information symmetrically as follows:

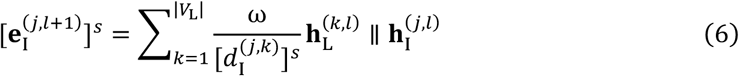

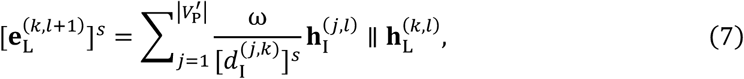

where ∥ is the concatenation operator, *l* is the number of the layer, and *s* is the exponent of the pairwise distance.

For different force fields, exponents of the distance are different. For example, *s* = 1 is used to model the electrostatic interaction, while *s* = 6 or 12 is used in the Lennard-Jones potential model to describe the attractive and repulsive energy. To capture different interaction patterns, in the EIRL, for a protein, multiple terms with different exponents are combined like an empirical SF as follows:

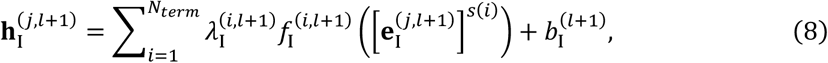

where 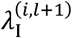 is a learnable weighting coefficient, 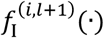 is an MLP, *N*_term_ is the number of terms, and *s*(*i*) is the function which determines the exponent of each term. *N*_*term*_ is a hyperparameter and set to 5 in this study. The *s*(*i*) is designed as follows:

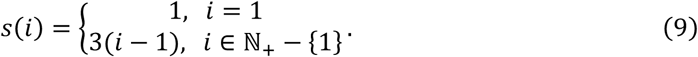

Similarly, the embeddings 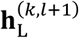 of atoms in a ligand are also updated but with different learnable weighting coefficients 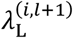 and parameters.

Two EIRLs are stacked and their learned residue/atom embeddings are concatenated. In the readout block, a global max pooling layer is used to aggregate all residue/atom embeddings into the embeddings of the protein/ligand. Then, the embeddings of the protein and the ligand are concatenated as the input of an MLP to predict p*K*_*d*_ value of the protein-ligand complex.

### 2.4. Feature importance analysis using integrated gradients

EGNA extracts multiple types of raw node features for proteins and ligands. To investigate the importance of these features, we apply a popular attribution method, *integrated gradients*, to each sample (i.e., complex) in CASF-2016 and CSAR-HiQ (Sundararajan, et al., 2017). Integrated gradients first design a baseline input, the size of which is the same to the raw feature matrix. In this study, the baselines are set to all-zero matrices. Then, the Riemann approximation of the integral of integrated gradient with respect to each input feature element is calculated. We denote the integrated gradients of a sample as **G** ∈ ℝ^|*V*|×*m*^, where |*V*| is the number of nodes in the graph and *m* is the length of feature vectors. The contribution of a specific feature type is calculated as follows:

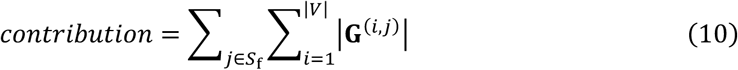

where *S*_f_ is the set of the indices of the features corresponding to this feature type. For example, only evolutionary features are used for proteins, and thus *S*_f_ = ℕ_+_ ∩ [1,30] for protein features. Similarly, the atom type corresponds to the 1^st^-9^th^ columns of **G** and thus *S*_f_ = ℕ_+_ ∩ [1,9] for it. Contribution of ligand features is the sum of that of atom type, aromaticity, in ring, hydrogen-bond acceptor, hybridization and partial charge.

## 3. Results and discussions

In this section, we compare our model EGNA with existing scoring functions on the CASF-2016 and CSAR-HiQ. Beyond the overall statistics, we further investigate whether the scaffold regions are beneficial for accurate binding affinity prediction.

### 3.1. Evaluation protocols

The goal of scoring functions is to predict the equilibrium dissociation constant (*K*_*d*_) of the protein-ligand complexes with known bound structures. The Gibbs free energy can be easily calculated when *K*_*d*_ is known. For convenience, p*K*_*d*_ (i.e., − log *K*_*d*_) is used as the target value and the unit of p*K*_*d*_ is Mol.

Three widely used metrics are selected for performance evaluation. Root mean squared error (RMSE), mean absolute value (MAE), and Pearson correlation coefficient (*R*) are defined as:

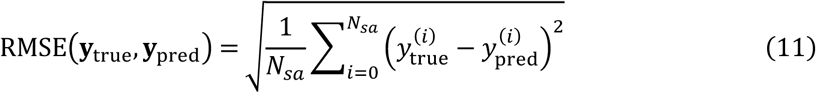

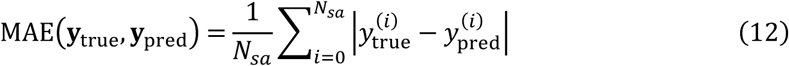

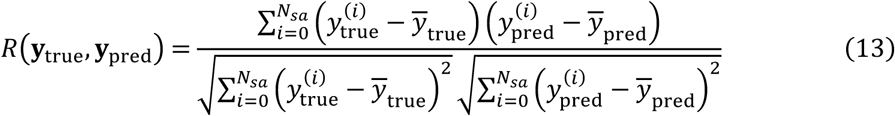

where 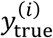 and 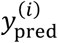 are the experimentally determined and predicted p*K*_*d*_ of the *i*^th^ sample in a dataset, respectively. *N*_*sa*_ is the number of samples in the dataset. Higher R, lower RMSE and lower MAE mean better performance.

The whole general set and 80% randomly selected samples of the refined set of PDBbind v.2016 are used as the training set. The remaining samples of the refined set are used as the validation set. The hyperparameters are determined by minimizing the RMSE on the validation set. The CASF-2016 and CSAR-HiQ are used as the independent testing sets.

### 3.2. Comparison with the state-of-the-art scoring functions

In recent years, various machine learning-based SFs are proposed for protein-ligand binding affinity prediction, including conventional machine learning (ML) models and deep learning (DL) models. We compare our method EGNA with these state-of-the-art SFs, which are briefly introduced in the Section D in the S1 Supporting Information (Jimenez, et al., 2018; Li, et al., 2015; Li, et al., 2021; Meng and Xia, 2021; Nguyen and Wei, 2019; Seo, et al., 2021; Stepniewska-Dziubinska, et al., 2018; Wojcikowski, et al., 2019; Zheng, et al., 2019). It should be noted that the unit of RMSE for AGL-Score and PerSpect is kcal/Mol, which is transformed to p*K*_*d*_ by dividing 1.3633 (Wang, et al., 2017).

As shown in Table 1, conventional ML-based SFs and DL-based SFs have comparable performance on CASF-2016. One reason for this phenomenon is that deep learning models usually require a huge volume of data for model training. Nevertheless, the number of protein-ligand complexes with known structures and binding affinities is still limited (i.e., <20k in PDBbind). In spite of the lack of annotated data, EGNA achieves the highest *R* and the lowest RMSE and MAE among all the methods. PerSpect is the second-best method, whose performance is close to that of our method EGNA. PerSpect is a ML-based SF and would heavily be dependent on the feature engineering; in contrast, EGNA is an end-to-end learning framework which only requires a combination of raw features of amino acids and atoms and is expected to have a better generalization ability. Our experiments also show that the *R* of EGNA is at least 2.2% higher than that of other DL-based SFs. A potential reason is that the intermolecular distance information is not fully exploited by other existing methods. However, in our model, the distance between any residues on the binding pocket and any atoms in the ligand are captured in EGNA. The relationship between intermolecular distance and residue/atom properties can be modeled more easily by the proposed empirical interaction representation layers.

**Table 1.**
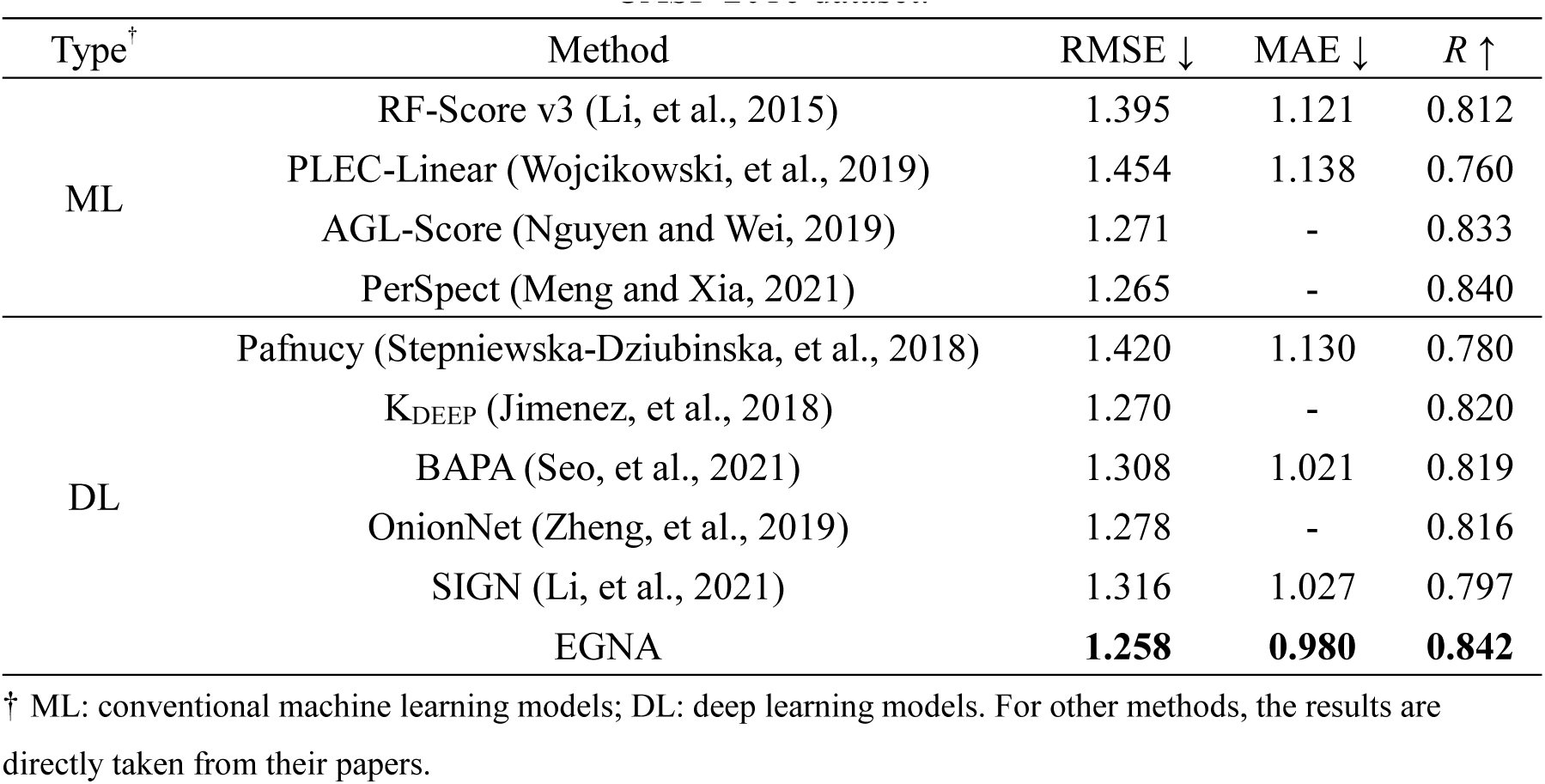
Comparison between the state-of-the-art machine learning-based SFs with EGNA on CASF-2016 dataset.

| Type<sup>†</sup> | Method | RMSE ↓ | MAE ↓ | $R$ ↑ |
| --- | --- | --- | --- | --- |
| ML | RF-Score v3 (Li, et al., 2015) | 1.395 | 1.121 | 0.812 |
|  | PLEC-Linear (Wojcikowski, et al., 2019) | 1.454 | 1.138 | 0.760 |
|  | AGL-Score (Nguyen and Wei, 2019) | 1.271 | - | 0.833 |
|  | PerSpect (Meng and Xia, 2021) | 1.265 | - | 0.840 |
| DL | Pafnucy (Stepniewska-Dziubinska, et al., 2018) | 1.420 | 1.130 | 0.780 |
|  | K<sub>DEEP</sub> (Jimenez, et al., 2018) | 1.270 | - | 0.820 |
|  | BAPA (Seo, et al., 2021) | 1.308 | 1.021 | 0.819 |
|  | OnionNet (Zheng, et al., 2019) | 1.278 | - | 0.816 |
|  | SIGN (Li, et al., 2021) | 1.316 | 1.027 | 0.797 |
|  | EGNA | **1.258** | **0.980** | **0.842** |

<sup>†</sup> ML: conventional machine learning models; DL: deep learning models. For other methods, the results are directly taken from their papers.

To evaluate the generalization ability, those methods which release their trained models or provide an online server are tested with EGNA on the nonoverlapping CASR-HiQ. The results summarized in Table 2 show that the performance of all methods degrades with different margins due to data distribution shift. EGNA still achieves the best RMSE, MAE, and *R* among all the methods. Compared with the second-best method, PLEC-Linear, the RMSE/MAE is decreased about 7.2%/4.2% and *R* is increased about 0.9% by our model EGNA. Compared to other DL-based SFs, the RMSE/MAE of EGNA decreases over 5.8%/5.3% and *R* of EGNA increases more than 2.2%. The results are statistically significant according to the statistical hypothesis test in (Meng and Kurgan, 2016), which is described in the Section E in the S1 Supporting Information in detail. The results on two datasets suggest that EGNA can effectively improve the accuracy of binding affinity prediction and yields high generalization performance.

**Table 2.** Comparison between the state-of-the-art machine learning-based SFs with EGNA on nonoverlapping CSAR-HiQ dataset.

| Type | Method<sup>†</sup> | RMSE ↓ | MAE ↓ | $R$ ↑ |
| --- | --- | --- | --- | --- |
| ML | RF-Score v3 | 1.917** | 1.433** | 0.532** |
|  | PLEC-Linear | 1.656** | 1.242** | 0.741* |
| DL | Pafnucy | 1.903** | 1.602** | 0.620** |
|  | K<sub>DEEP</sub> | 1.630** | 1.257** | 0.710** |
|  | BAPA | 1.700** | 1.346** | 0.669** |
|  | OnionNet | 1.714** | 1.319** | 0.661** |
|  | EGNA | **1.536** | **1.190** | **0.750** |

<sup>†</sup> The results are obtained by running their standalone software without retraining or submitting jobs to their online servers.
\* $p$ -value of t-test is $< 0.05$ . \*\* $p$ -value of t-test is $< 0.01$ .

### 3.3. Residues in scaffold regions contribute to binding affinity prediction for EGNA

When constructing the protein and interaction graphs, residues in the protein are divided into two types according to the distance between their *C*_*β*_ and the closest atom in the ligand. The residues belong to 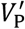 are considered as the members of binding pockets while those belonging to 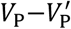 are considered as the members of scaffold regions. Thus, the range of the distance between the residues in the scaffold regions and their closest atoms is (10, *d*_*t*_] according to the definitions in Section 2.2.1. Intuitively, the binding affinity is mostly related to the residues in the binding pockets. In a previous GNN-based method, SIGN (Li, et al., 2021), only binding pockets are taken into account. However, if *d*_*t*_ = 20Å, residues in the scaffold region are usually one- or two-hop neighbors of those in the binding pockets in graphs. Thus, it is a reasonable assumption that our model EGNA can benefit from larger protein graphs, which are composed of residues from both binding pockets and scaffold regions.

To validate this assumption, we additionally set *d*_*t*_ to 10Å or 30Å to investigate the impacts of the scaffold region in proteins. Our model EGNA is retrained accordingly and the results are shown in Fig 3. When *d*_*t*_ = 10Å, there is no residue in scaffold region and the retrained model EGNA achieves the lowest *R* and highest RMSE on both datasets. The results demonstrate that scaffold regions can certainly provide useful information related to the binding affinity. When *d*_*t*_ = 30Å, our retrained model achieves a slightly higher *R* (0.849 vs 0.842) but also a higher RMSE (1.261 vs 1.258) on CASF-2016, and is comparable to the original model with *d*_*t*_ = 20Å. However, its RMSE (1.641 vs 1.536) and *R* (0.705 vs 0.750) are both inferior to the original model on CSAR-HiQ. A potential reason for the performance deterioration is that a larger scaffold region may introduce noise and lead to overfitting. Overall, our results suggest that choosing an appropriate size of the scaffold region is important for the protein representation and can improve the accuracy of protein-ligand binding affinity prediction.

**Fig 3.**
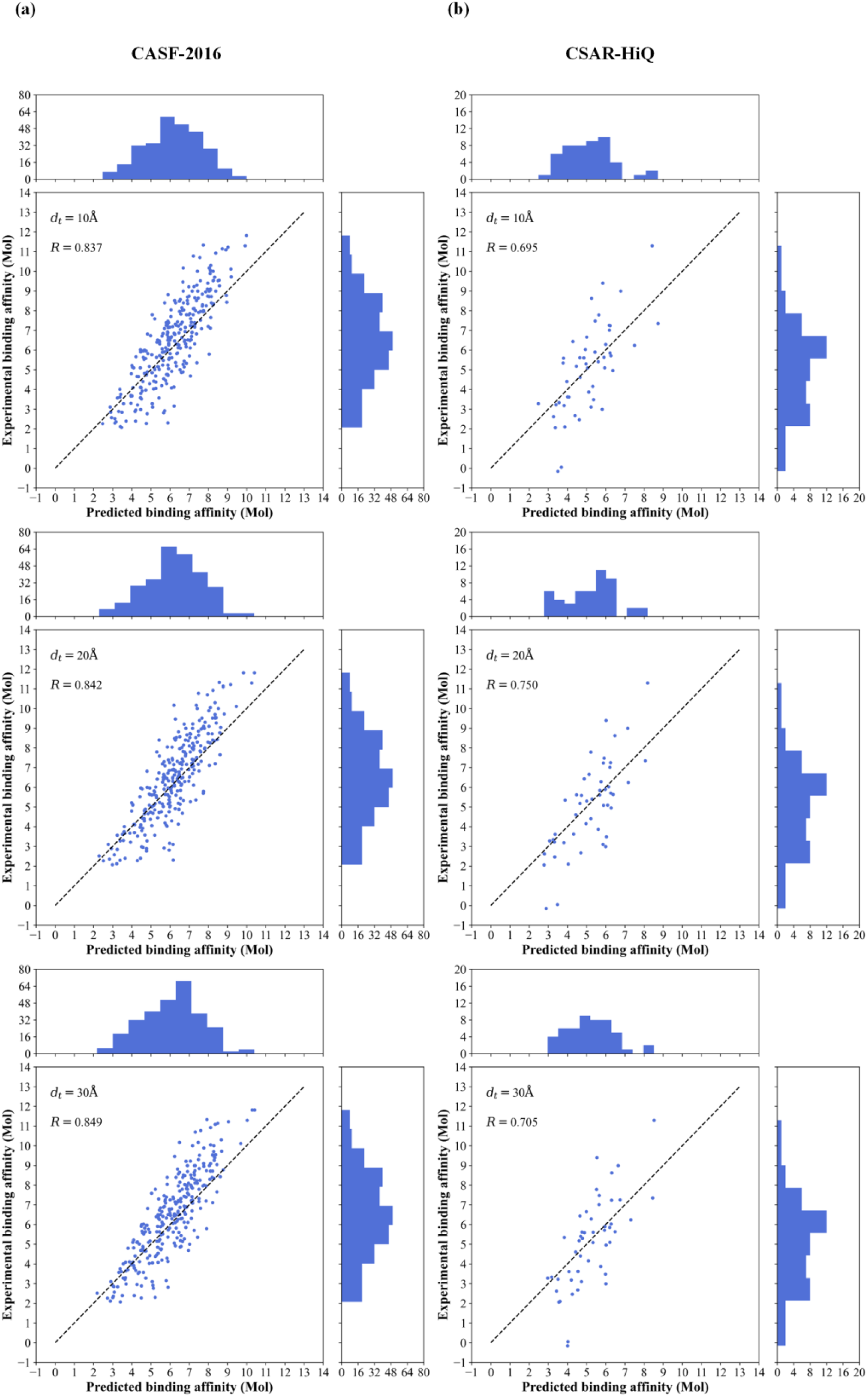
Comparison of predicted binding affinity and experimental binding affinity under different sizes of scaffold regions on (a) CASF-2016 and (b) CSAR-HiQ. When *d*_*t*_ = 10/20/30Å, the RMSE is 1.278/1.258/1.261Å on CASF-2016 and 1.643/1.536/1.641Å on CSAR-HiQ.

### 3.4. Empirical interaction representation layers improve the binding affinity prediction

Empirical interaction representation layers (EIRLs) are used to represent the bimolecular interaction in a way similar to empirical SFs. Multiple pseudo energy terms in EIRLs are weighted and summed in Eq.8. EIRLs are influenced by the number of terms and the exponent of the distance. If we set *N*_*term*_ = 1 and *s*(*i*) = 1, EIRLs are degenerated into standard graph convolutional layers (GCLs). To demonstrate the advantages of EIRLs in EGNA, we substitute the EIRLs with GCLs and retrain EGNA as EGNA-GCL.

As shown in Fig 4, the *R* of EGNA-GCL is 1.6% and 4.1% lower than the original EGNA on CASF-2016 and CSAR-HiQ, respectively. The RMSE of EGNA-GCL also increases 6.4% and 7.7% over the original EGNA, respectively. The results illustrate that EIRLs are more powerful than the standard GCLs in EGNA. However, it is still possible that the improvement brought by EIRLs is due to more learnable parameters. To test this concern, we set *N*_*term*_ = 5 to keep the number of parameters unchanged and the variants are named as multi-channel graph convolutional layers. The results in Fig 4 show that the *R* and RMSE of EGNA with multi-channel GCLs (EGNA-MGCL) are nearly the same to those of standard GCLs on CASF-2016 and inferior to EIRLs. When multi-channel GCLs are evaluated on CSAR-HiQ, the *R* is increased and close to that of EIRLs. However, the RMSE of multi-channel GCLs is still 3.8% higher than that of EIRLs. These results indicate the overall performance of EIRLs are better than standard and multi-channel GCLs on the protein-ligand interaction representation.

**Fig 4.**
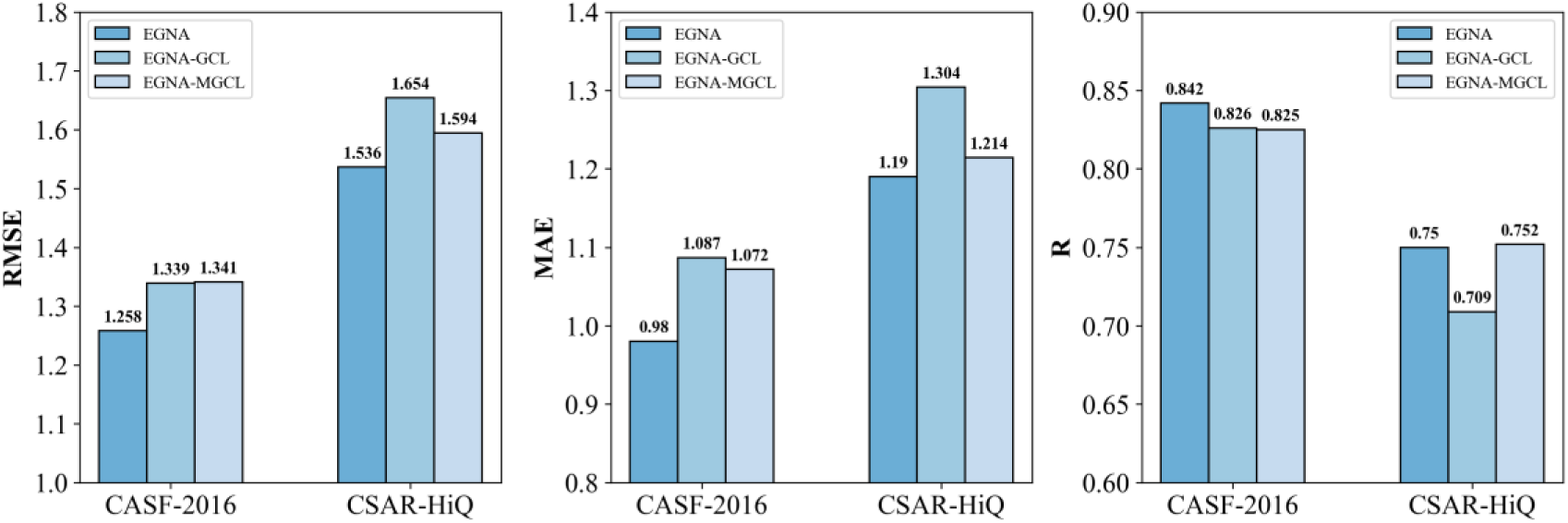
Comparison of EGNA, EGNA-GCL and EGNA-MGCL on CASF-2016 and CSAR-HiQ. The *p*-values of t-test show that the difference between EGNA, EGNA-GCL, and EGNA-MGCL is significant (*p*-value < 0.05) except the *R* of EGNA and EGNA-MGCL on CSAR-HiQ (*p*-value=0.22).

To investigate the impacts of different pseudo energy terms, we extract the learnable weighting coefficients λ of EIRLs (ref. Eq.9) and multi-channel GCLs (*s*(*i*) = 1) from the trained models. The absolute values of 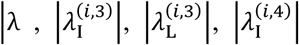,and 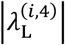, for the two layers (EIRLs and multi-channel GCLs) are shown in Fig 5. For multi-channel GCLs, λ of different terms are close in absolute value as expected. The results demonstrate that these terms contribute nearly equally to the learned atom/residue embeddings. The reason is that the input of each term is exactly the same and all trainable parameters are initialized similarly.

**Fig 5.**
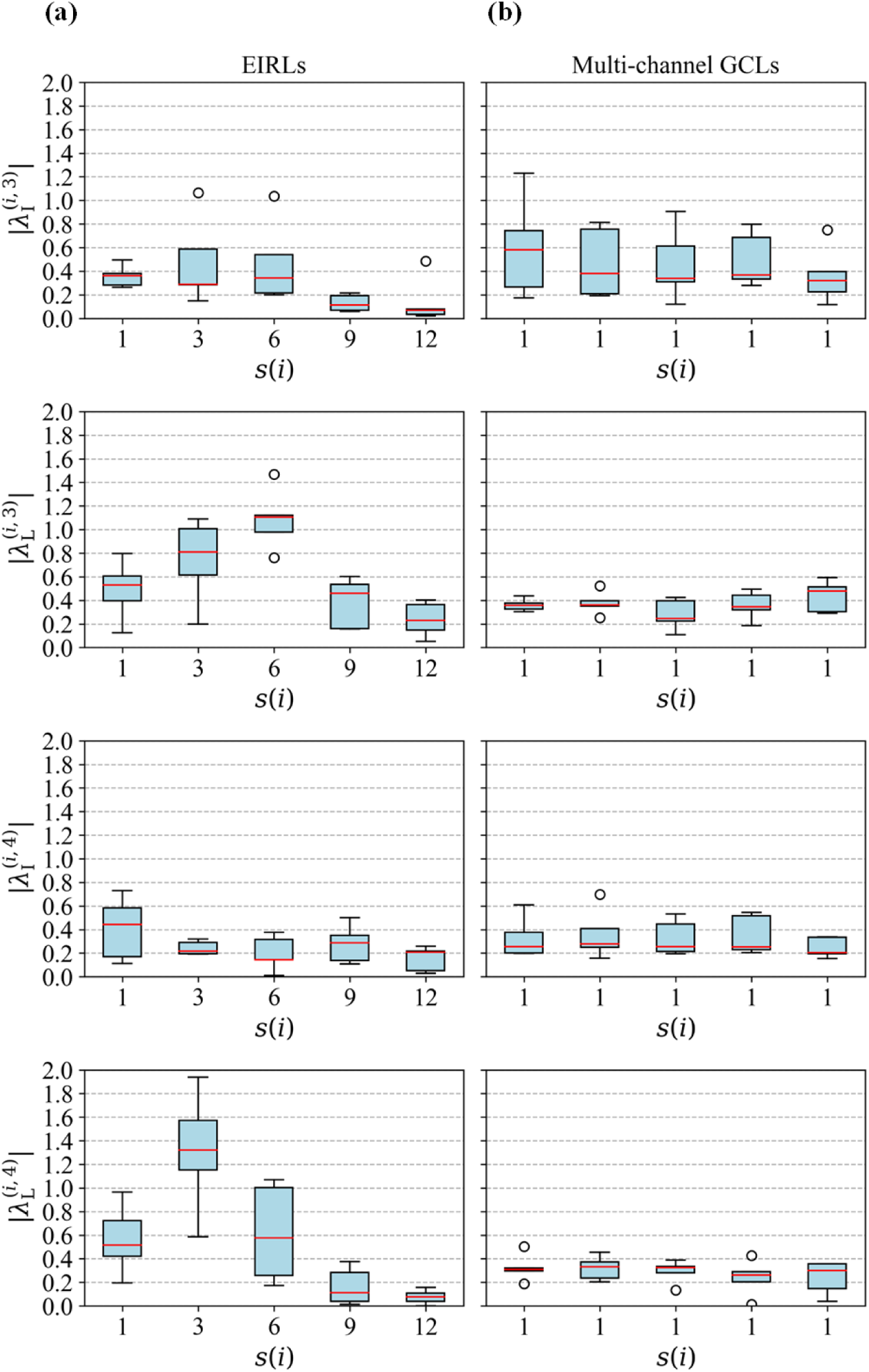
Comparison between the learnable weighting coefficients of (a) empirical interaction representation layers (ref. Eq.9) and (b) multi-channel graph convolutional layers (*s*(*i*) = 1). Considering that the predicted binding affinity is an average of the outputs from five independently trained models, the weighting coefficients of all five models are summarized.

Differently, 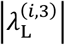 and 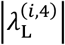 of different terms in EIRLs vary considerably. 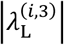 achieves its maximum and minimum performance at *s*(*i*) = 6 and *s*(*i*) = 12, respectively. 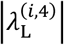 is similar to 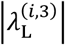 but achieves its maximum at *s*(*i*) = 3. Obviously, those terms with *s*(*i*) = 3 or 6 are more influential in the output than those with *s*(*i*) = 12. Although the relationship between different terms cannot be figured out as easily as a Lennard-Jones potential model due to the nonlinear transformation in high-dimensional space, it is clear that EIRLs improve the representation capability of neural networks by introducing the pairwise distance with diverse exponents.

### 3.5. Raw node features of proteins contribute more to the binding affinity

In our model EGNA, multiple types of raw node features are used to represent residues in proteins and heavy atoms in ligands. Thus, we further dig into the trained model to investigate the contribution of individual raw node features using integrated gradients.

The mean and standard deviation of the contribution of each feature type are calculated as Eq.10 and shown in Fig 6. The results demonstrate that the output of our model EGNA is generally more influenced by protein features than ligand features. It also should be noted that the standard deviations of both features are relatively large, indicating that the contributions are highly dependent on the samples. For example, protein features contribute less to the predicted p*K*_*d*_ values in CASF-2016 than those in CSAR-HiQ. Overall, protein and ligand features are both necessary for EGNA but protein information are more related to binding affinity prediction.

**Fig 6.**
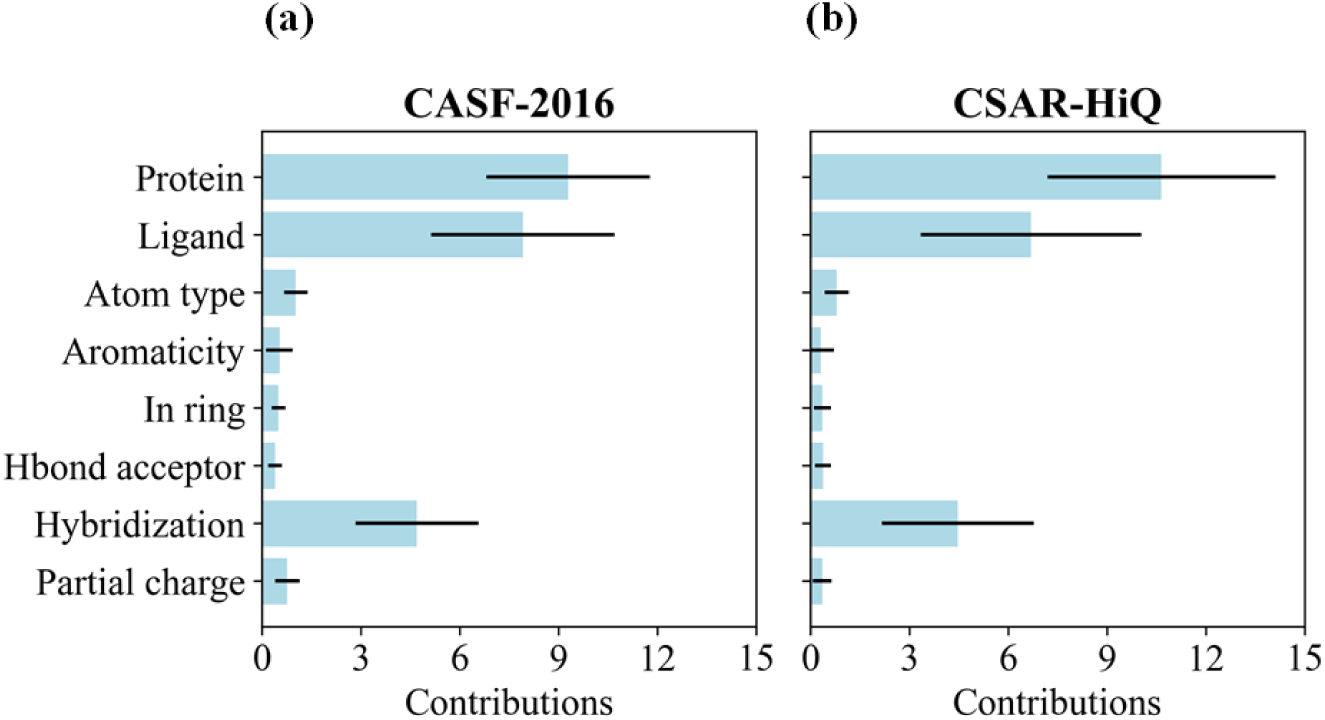
Contributions of raw protein and ligand node features using integrated gradients on (a) CASF-2016 and (b) CSAR-HiQ. Protein node features are sequence profiles derived from HHblits. Ligand node features are composed of the atom type, aromaticity, in ring, hydrogen-bond acceptor, hybridization and partial charge.

Contrary to protein features, there are multiple types of atom properties used as the ligand features. The contribution of atom hybridization is much larger than other atom properties. A potential reason is that atom hybridization state provides more detailed description about the local structures and bonds of the ligands. Atom type is the second most important factor. Considering that element-specific pairwise distance alone has been proved an effective feature for protein-ligand binding affinity prediction in RF-Score (Ballester and Mitchell, 2010), the importance of atom type is consistent with our expectations. Partial charge is correlated to the electrostatic potential energy and its contribution is slightly higher than the rest of features on CASF-2016 but shows no difference on CSAR-HiQ. Other atom features also contribute to the final prediction but their impacts are relatively limited.

## 4. Conclusions

In this study, we propose a novel approach EGNA based on empirical graph neural networks for accurate protein-ligand binding affinity prediction. The interaction between proteins and ligands are modelled by a dense graph simulating an empirical scoring function. The empirical interaction representation layer can capture the finer-grained interaction patterns by fully exploiting the intermolecular distance information, which result in better prediction performance. We also find that scaffold regions of proteins provide extra useful information for binding affinity prediction other than the binding pockets. The learnable weighting coefficients of EIRLs also enable us to reveal different contributions of raw node features.

In the experiments, we observed that the performance of all ML-based SFs trained on PDBbind degraded on CSAR-HiQ. This phenomenon illustrates that ML-based SFs are still restricted by insufficient and low-quality training data. In the future, if more annotated data are available, the performance of ML-based SFs, especially DL-based SFs, can be further improved. Considering that the structures of many unannotated complexes are known, unsupervised or semi-supervised machine learning models may also be introduced to enhance the prediction model. Moreover, it is also possible to extend our model to protein-protein or protein-nucleotide interaction prediction.

